# ExpansionHunter Denovo: A computational method for locating known and novel repeat expansions in short-read sequencing data

**DOI:** 10.1101/863035

**Authors:** Egor Dolzhenko, Mark F. Bennett, Phillip A. Richmond, Brett Trost, Sai Chen, Joke J.F.A. van Vugt, Charlotte Nguyen, Giuseppe Narzisi, Vladimir G. Gainullin, Andrew Gross, Bryan Lajoie, Ryan J. Taft, Wyeth W. Wasserman, Stephen W. Scherer, Jan H. Veldink, David R. Bentley, R K.C. Yuen, Melanie Bahlo, Michael A. Eberle

## Abstract

Expansions of short tandem repeats are responsible for over 40 monogenic disorders, and undoubtedly many more pathogenic repeat expansions (REs) remain to be discovered. Existing methods for detecting REs in short-read sequencing data require predefined repeat catalogs. However recent discoveries have emphasized the need for detection methods that do not require candidate repeats to be specified in advance. To address this need, we introduce ExpansionHunter Denovo, an efficient catalog-free method for genome-wide detection of REs. Analysis of real and simulated data shows that our method can identify large expansions of 41 out of 44 pathogenic repeats, including nine recently reported non-reference REs not discoverable via existing methods.

ExpansionHunter Denovo is freely available at https://github.com/Illumina/ExpansionHunterDenovo

## Background

High-throughput whole-genome sequencing (WGS) has experienced rapid reductions in per-genome costs over the past ten years (Muir et al. 2016) driving population-level sequencing projects and precision medicine initiatives at an unprecedented scale (Erikson et al. 2016; Telenti et al. 2016; Gudbjartsson et al. 2015; Nagasaki et al. 2015; 1000 Genomes Project Consortium et al. 2015; Consortium and Project MinE ALS Sequencing Consortium 2018). The availability of large sequencing datasets now allows researchers to perform comprehensive genome-wide searches for disease-associated variants. The primary limitations of these studies are the completeness of the reference genome and the ability to identify putative causal variations against the reference background. A wide variety of software tools can identify variations relative to the reference genome such as single nucleotide variants (SNVs) and short (1-50 bp) insertions and deletions (indels) (McKenna et al. 2010; Raczy et al. 2013; Rimmer et al. 2014; Poplin et al. 2016; Garrison and Marth 2012; Poplin et al. 2018), copy number variants (CNVs) (Roller et al. 2016; Abyzov et al. 2011) and structural variants (SVs) (Chen et al. 2016; Layer et al. 2014; Abyzov et al. 2011). A common feature of these variant callers is their reliance on sequence reads that at least partially align to the reference genome. However, because some variants include large amounts of inserted sequence relative to the reference, methods that can analyze reads that do not align to the reference are also needed.

A particularly important category of variants that involve long insertions are repeat expansions (REs) such as the expansions in *C9orf72* repeat associated with amyotrophic lateral sclerosis (ALS). This repeat consists of three copies of CCGGGG motif in the reference (18 bp total) whereas the pathogenic mutations are comprised of at least 30 copies of the motif (180 bp total) and may reach into thousands of bases (DeJesus-Hernandez et al. 2011; Renton et al. 2011). REs are known to be responsible for dozens of monogenic disorders (La Spada and Paul Taylor 2010; Hannan 2018).

Several recently-developed tools can detect REs longer than the standard short read sequencing read length of 150 bp (Tang et al. 2017; Tankard et al. 2018; Dolzhenko et al. 2017; Dashnow et al. 2018; Dolzhenko et al. 2019; Mousavi et al. 2019). These tools have all been demonstrated to be capable of accurately detecting pathogenic expansions of simple short tandem repeats (STRs). However, recent discoveries have shown that many pathogenic repeats have complex structure and hence require more flexible methods. For instance: (a) REs causing spinocerebellar ataxia types 31 and 37, familial adult myoclonic epilepsy types 1, 2, 3, and 4, and Baratela-Scott syndrome (Sato et al. 2009; Seixas et al. 2017; Ishiura et al. 2018; Corbett et al. 2019; Florian et al. 2019; Yeetong et al. 2019; LaCroix et al. 2019) occur within an inserted sequence relative to the reference; (b) expanded repeats recently shown to cause spinocerebellar ataxia, familial adult myoclonic epilepsy, and cerebellar ataxia with neuropathy and bilateral vestibular areflexia syndrome have different composition relative to the reference STR (Sato et al. 2009; Seixas et al. 2017; Ishiura et al. 2018; Corbett et al. 2019; Florian et al. 2019; Yeetong et al. 2019); (c) Unverricht-Lundborg disease, a type of progressive myoclonus epilepsy, is caused by an expansion of a dodecamer (12-mer) repeat (Lalioti et al. 1997). None of the existing methods are capable of discovering all of these REs.

We have developed ExpansionHunter Denovo (EHdn), a novel method for performing genome-wide search for expanded repeats, to address the limitations of the existing approaches. EHdn scans the existing alignments of short reads from one or many sequencing libraries, including the unaligned and misaligned reads, to identify approximate locations of long repeats and their nucleotide composition. This method (a) does not require prior knowledge of the genomic coordinates of the REs, (b) can detect nucleotide composition changes within the expanded repeats, and (c) is applicable to both short and long motifs. EHdn is computationally efficient because it does not re-align reads. Depending on the sensitivity settings, EHdn can analyze a single 30-40x WGS sample in about 30 minutes to 2 hours on a single CPU thread.

In this study, we demonstrate that EHdn can be used to rediscover the REs associated with fragile X syndrome (FXS), Friedreich Ataxia (FRDA), Myotonic Dystrophy type 1 (DM1), and Huntington’s disease (HD) using case-control analysis to compare a small number of affected individuals (N=14-35) to control samples (N=150). We also show that REs in individual samples can be identified using outlier analysis. We then characterize large (longer than the read length) repeats in our control cohort to get a sense of baseline variability of these long repeats. Finally, we demonstrate the capabilities of our method by analyzing simulated expansions of various classes of tandem repeats known to play an important role in human disease. Taken together, our findings demonstrate that EHdn is a robust tool for identifying previously-unknown pathogenic repeat expansions in both cohort and single-sample outlier analysis, capable of identifying a new, previously inaccessible class of REs.

## Results

### ExpansionHunter Denovo

#### Overview

The length of disease-causing REs tends to exceed the read length of modern short-read sequencing technologies (Ashley 2016). Thus pathogenic expansions of many repeats can be detected by locating reads that are completely contained inside the repeats. As in our previous work (Dolzhenko et al. 2017, 2019), we call these reads in-repeat reads (IRRs). We implemented a method, ExpansionHunter Denovo (EHdn), for performing a genome-wide search for IRRs in BAM/CRAM files containing read alignments. EHdn computes genome-wide STR profiles containing locations and counts of all identified IRRs. Subsequent comparisons of STR profiles across multiple samples can reveal the locations of the expanded repeats.

#### Genome-wide STR profiles

Genome-wide STR profiles computed by EHdn contain information about two types of IRRs: anchored IRRs and paired IRRs. Anchored IRRs are IRRs whose mates are confidently aligned to the genomic sequence adjacent to the repeat (Methods). Paired IRRs are read pairs where both mates are IRRs with the same repeat motif. Repeats exceeding the read length generate anchored IRRs (Figure 1, middle panel). Repeats that are longer than the fragment length of the DNA library produce paired IRRs in addition to anchored IRRs (Figure 1, right panel). The genomic coordinates where the anchored reads align correspond to the approximate locations of loci harboring REs and the number of IRRs is indicative of the overall RE length.

**Figure 1.**
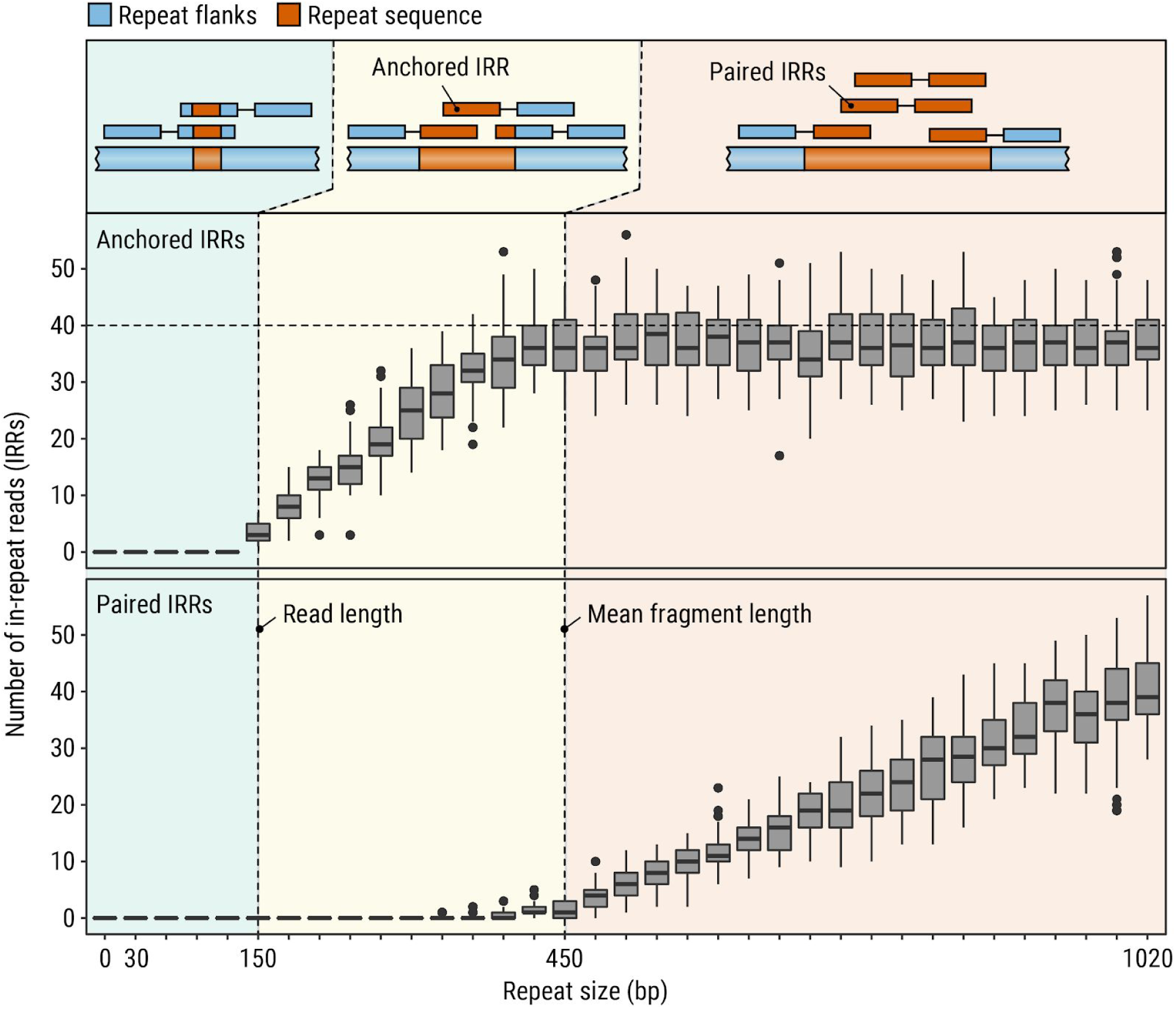
Diagram illustrating the types and counts of reads generated by simulating repeats of different lengths. When the repeat is shorter than the read length (left panels), there are no IRRs associated with the repeat. When a repeat is longer than the read length but shorter than the fragment length (middle panels), anchored IRRs but no paired IRRs are present. As the repeat length approaches and exceeds the fragment length (right panels), paired IRRs are generated in addition to anchored IRRs.

The information about anchored IRRs is summarized in an STR profile for each repeat motif (e.g. CCG) by listing regions containing anchored IRRs in close proximity to each other together with the total number of anchored IRRs identified there (Figure 2, middle). Note that the mapping positions of anchored IRRs correspond to the positions of anchor reads; mapping positions of IRRs themselves are not used because their alignments are often unreliable. Contrary to anchored IRRs, the origin of paired IRRs cannot be determined if a genome contains multiple long repeats with the same motif. Due to this, STR profiles only contain the overall count of paired IRRs for each repeat motif.

**Figure 2.**
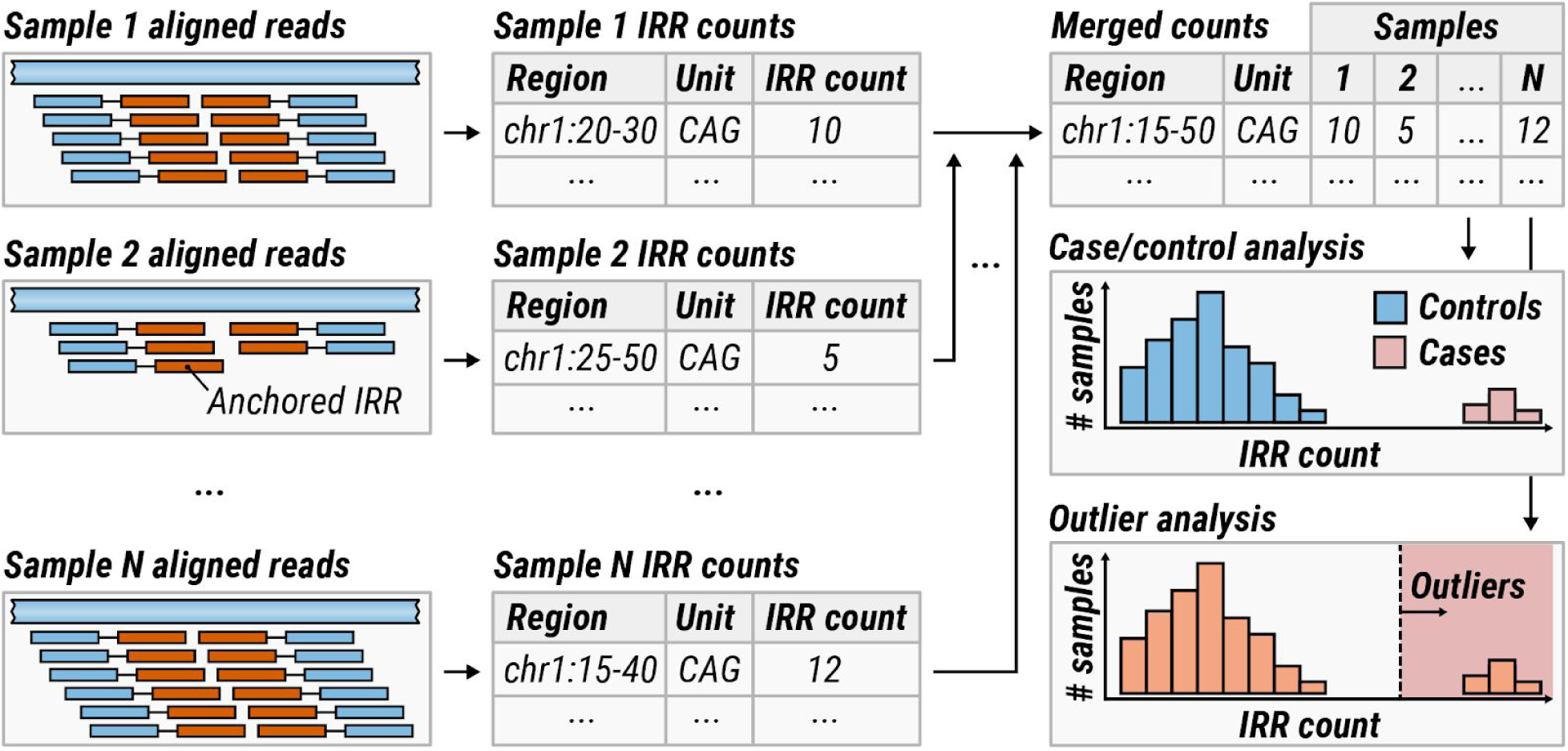
(Left) A search for anchored IRRs is performed across all aligned reads. (Middle) The IRR counts are summarized into STR profiles. (Right) The resulting STR profiles are merged across all samples. If the dataset can be partitioned into cases and controls, IRR counts in these groups are compared for each locus. Alternatively, if no such partition is possible, an outlier analysis is performed.

#### Comparing STR profiles across multiple samples

To compare STR profiles across multiple samples, the profiles must first be merged together. During this process, nearby anchored IRR regions are merged across multiple samples and the associated counts are depth-normalized and tabulated for each sample (Figure 2, right; Methods). The total counts of paired IRRs are also normalized and tabulated for each sample. The resulting per-sample counts can be compared in two ways: If the samples can be partitioned into cases and controls where a significant subset of cases is hypothesized to contain expansions of the same repeat then a case/control analysis can be performed using a Wilcoxon rank-sum test (Methods). Alternatively, if no enrichment for any specific expansion is expected, an outlier analysis (Methods) can be used to flag repeats that are expanded in a small subgroup of cases compared to the rest of the dataset. Case-control and outlier analyses can be performed on either anchored IRRs or paired IRRs, which we call locus and motif methods, respectively (Methods). Thus, the locus method can reveal locations of repeat expansions while the motif method can reveal the overall enrichment for long repeats with a given motif.

#### Baseline simulations

To demonstrate the baseline expectation of how the numbers of anchored and paired IRRs vary with repeat length, we simulated 2×150 bp reads at 20x coverage with 450 bp mean fragment length for the repeat associated with Huntington’s disease and varied the repeat length from 0 to 340 CAG repeats (0 to 1,020 bp; Supplemental Information). No IRRs occur when the repeat was shorter than the read length (Figure 1, left panel). When the repeat is longer than the read length but shorter than the fragment length (Figure 1, middle panel), the number of anchored IRRs increases proportionally to the length of the repeat. As the length of the repeat approaches and exceeds the mean fragment length (Figure 1, right panel), the number of paired IRRs increases linearly with the length of the repeat. Because anchored IRRs require one of the reads to “anchor” outside of the repeat region, the number of anchored IRRs is limited by the fragment length and remains constant as the repeat grows beyond the mean fragment length. It is important to note that real sequence data may introduce additional challenges compared to the simulated data. For example, sequence quality in low complexity regions or interruptions in the repeat may impact the ability to identify some IRRs.

### Analysis of sequencing data

#### Detection of expanded repeats in case-control studies

Given a sufficient number of samples with the same phenotype, pathogenic REs may be identified by searching for regions with significantly longer repeats in cases compared to controls (see Figure 2). To demonstrate the feasibility of such analyses, we analyzed 91 Coriell samples with experimentally-confirmed expansions in repeats associated with Friedreich’s ataxia (FRDA; N=25), Myotonic Dystrophy type 1 (DM1; N=17), Huntington disease (HD; N=14), and fragile X syndrome (FXS; N=35). This dataset has been previously used to benchmark the performance of existing methods (Tankard et al. 2018; Dolzhenko et al. 2017; Mousavi et al. 2019).

The pathogenic cutoffs for FRDA, DM1, and FXS repeats are greater than the read length, so our analysis of simulated data suggests that anchored IRRs are likely to be present in each sample with one of these expansions (Figure 1). The pathogenic cutoff for the HD repeat (120 bp) is less than the read length (150 bp) used in this study, so a subset of samples with Huntington’s disease may not contain relevant IRRs making this expansion harder to detect, however it is detectable with existing methods (Dolzhenko et al. 2017, 2019; Tang et al. 2017; Mousavi et al. 2019)

We separately compared samples with expansions in *FXN* (FRDA), *DMPK* (DM1), *HTT* (HD), or *FMR1* (FXS) genes (cases) against a control cohort of 150 unrelated Coriell samples of African, European, and East Asian ancestry (Illumina n.d.). Each case-control comparison revealed a clear enrichment of anchored IRRs from the corresponding repeat region (Figure 3). This analysis demonstrated that ExpansionHunter Denovo can re-identify known pathogenic repeat expansions without prior knowledge of the location or repeat motif when the pathogenic repeat length is longer than (or nearly the size of) the read length and the repeat is highly penetrant.

**Figure 3:**
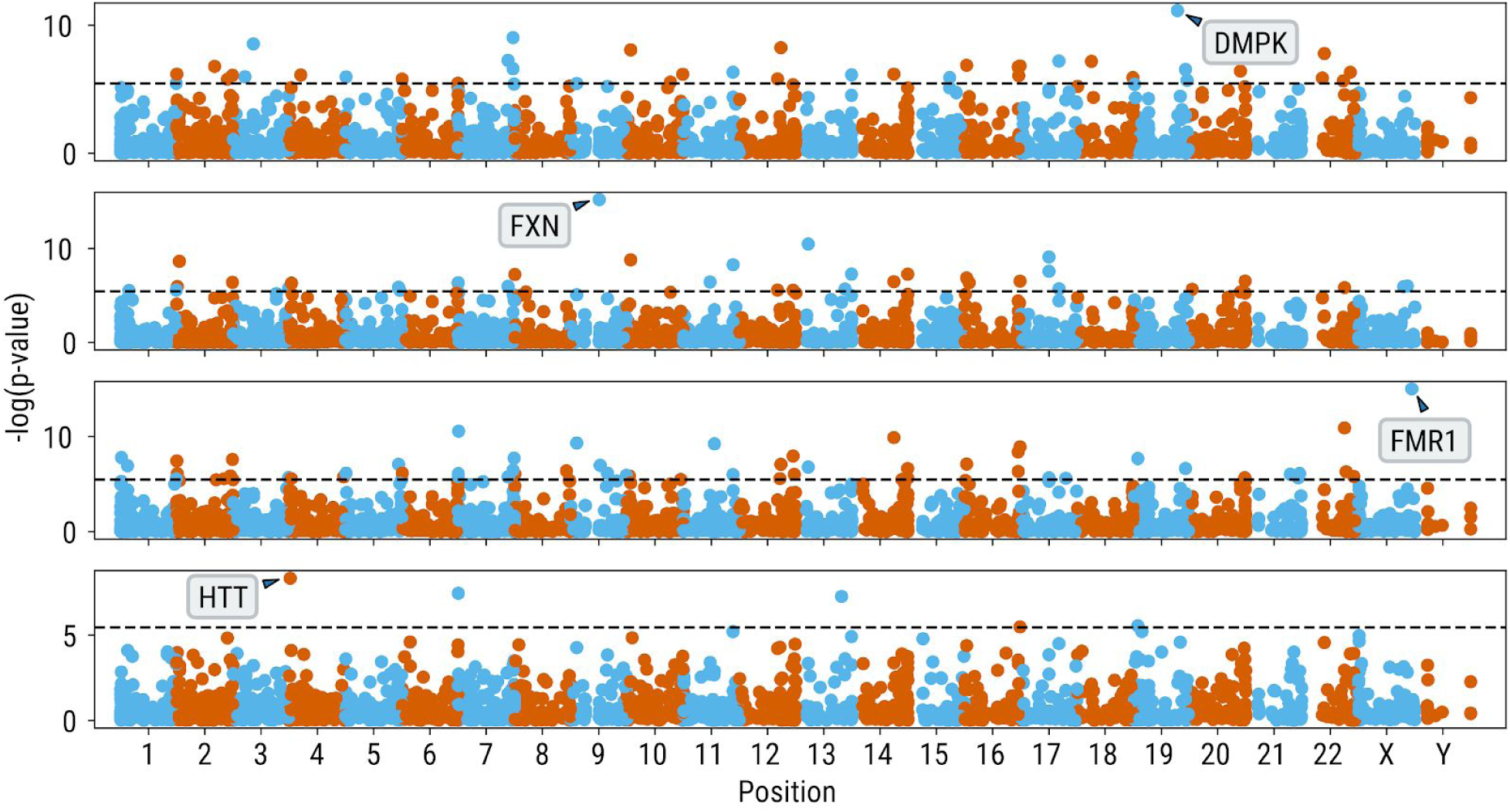
Genome-wide analysis of anchored IRRs comparing cases with known pathogenic expansions in *DMPK, FXN, FMR1* and *HTT* genes (top to bottom) to 150 controls.

#### Detection of expanded repeats in mixed sample cohorts

In many discovery projects it can be difficult to isolate patients that harbor the same repeat expansion based on the phenotype alone. For instance, the repeat expansion in the *C9orf72* gene is present in fewer than 10% of ALS patients and many ataxias can be caused by expansions of a variety of repeats. Such problems call for methods that are suitable for heterogeneous disease cohorts.

To solve this problem, we follow the approach taken previously by others and compare each case sample against the control cohort to identify outliers (Dashnow et al. 2018; Tankard et al. 2018) (Methods). To demonstrate the efficacy of this approach, we combined each sample from the pool of samples with expansions in *FXN, DMPK, HTT*, and *FMR1* genes with 150 controls to generate a total of 91 datasets, each containing 151 samples. We then performed an outlier analysis on the counts of anchored IRRs (Methods) in each dataset.

In 77% of the datasets, the expanded repeat ranked within the top 10 repeats based on the outlier score (Figure 4). This number increased to 82% when the analysis was restricted to short motifs between 2 and 6 bp. EHdn performed well for *DMPK* and *FXN* repeats, identifying these REs within the top 10 ranks for 41 out of 42 cases. The *FMR1* expansion was only ranked in the top 10 for 22 out of 35 cases known to have the expansion. This result is consistent with a previous comparison, which found this locus had poorest performance across all RE detection tools (Tankard et al. 2018). The performance for the *HTT* repeat is surprisingly good considering that EHdn was not designed to detect REs shorter than the read length.

**Figure 4:**
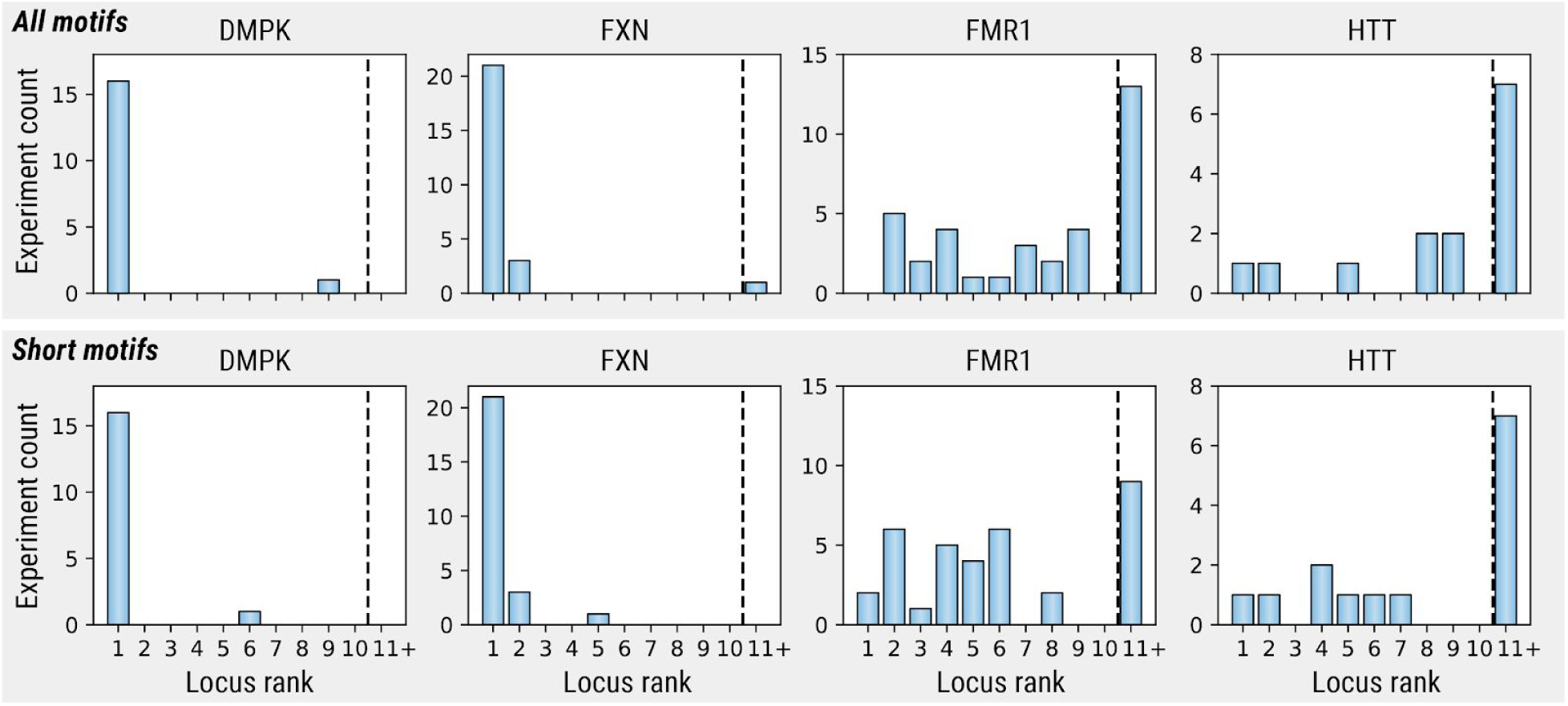
Ranking of known expansions based on the outlier score computed for anchored IRRs. Each rank originates from a genome-wide analysis of a dataset consisting of a single sample with a known expansion and 150 controls. (Top row) Ranks for all identified motifs. (Bottom row) Ranks for motifs 6bp and shorter.

#### The landscape of long repeats within a control population

To explore the landscape of large repeats in the general population, we applied EHdn to 150 unrelated Coriell samples of African, European, and East Asian ancestry (Illumina n.d.). To limit this analysis to higher confidence repeats we considered loci where EHdn identified at least five anchored IRRs and motifs supported by at least five paired IRRs in a single sample. Altogether, EHdn identified 1,572 unique motifs spanning between two and 20 bp, 94% of which were longer than 6 bp. Of these, 19% were found in at least half of the samples and 23% were found in just one sample. On average, each person had 658 loci with long repeats. As expected, the telomeric motif AACCCT is particularly abundant. It was found in about ∼23,000 IRRs per sample. Similarly, the centromeric motif AATGG was found in ∼5,000 IRRs per sample.

### Exploring the limitations of catalog-based RE detection methods

To evaluate the limitations of catalog-based approaches (Dolzhenko et al. 2017; Tang et al. 2017; Tankard et al. 2018; Dashnow et al. 2018; Mousavi et al. 2019), we curated a set of 53 pathogenic or potentially pathogenic repeats (Table S1) and checked if they were present in two commonly-used catalogs: (a) STRs with up to 6bp motifs from the UCSC genome browser simple repeats track (Benson 1999; Kent et al. 2002) utilized by STRetch and exSTRa and (b) the GangSTR catalog. Nine of the pathogenic repeats are not present in the reference genome and hence are absent from both catalogs. Out of the remaining 44 loci, 22 loci are present in both catalogs, 12 are missing from the GangSTR catalog and present in the UCSC catalog, five are missing from the UCSC catalog and present in the the GangSTR catalog, and five are missing from both catalogs (Figure S1). While it is possible to update the catalogs to include these known pathogenic repeats, the number of missing potentially-pathogenic REs remains unknown.

To demonstrate that EHdn offers similar performance to catalog-based methods on expansions exceeding the read length, we simulated expansions of the 35 non-degenerate STRs present in the reference (Table S1). We focused our comparisons on STRetch because this method was specifically designed to search for novel expansions using a genome-wide catalog and because it was shown to have similar performance to other existing methods (Tankard et al. 2018). Our simulations show that EHdn ranks 31out of 35 pathogenic repeats in the top 10 (Table S3-S6).

STRetch prioritizes 26 out of 29 repeats in the top 10 and the six remaining repeats are missing from its catalog. One of REs detected by EHdn and missed by STRetch is the pathogenic *CSTB* repeat with a motif length of 12bp. This is because STRetch is limited to detection of motifs with length up to six base pairs. To further highlight this strength of EHdn, we confirmed that it can detect other REs with long motifs (Supplemental Information).

Some recently-discovered REs are composed of motif not present the reference genome. One such example is the recently-discovered repeat expansion of non-reference motif AAGGG causing cerebellar ataxia with neuropathy and bilateral vestibular areflexia syndrome (CANVAS) (Cortese et al. 2019; Rafehi et al. 2019). Rafehi et al (Rafehi et al. 2019) demonstrated that EHdn is the only computational method capable of discovering this expansion. To further benchmark EHdn’s ability to detect REs with complex structure we simulated nine complex REs with non-reference motifs known to cause disease (Figure S3). For eight out of nine REs, including a simulated version of the CANVAS expansion, EHdn was able to detect one or both of the expanded repeats in each locus (Table S7).

## Discussion

Here, we introduced a new software tool, EHdn, that can identify novel REs using high-throughput WGS data. We tested EHdn by comparing samples with known REs against a control group of 150 diverse individuals and performed simulation studies across a range of pathogenic or potentially pathogenic REs. These analyses show that EHdn offers comparable performance to targeted methods on known pathogenic repeats while also being able to detect repeats absent from existing catalogs.

Recent discoveries have highlighted the importance of complex pathogenic repeat expansions involving non-reference insertions (Sato et al. 2009; Seixas et al. 2017; Ishiura et al. 2018; Corbett et al. 2019; Florian et al. 2019; Yeetong et al. 2019). EHdn is currently the only method capable of discovering these expansions from BAM or CRAM files without the need for re-alignment of the supporting reads. Additionally, we anticipate that EHdn can replace existing more manual and less computationally efficient discovery pipelines, such as the TRhist-based pipeline (Ishiura et al. 2019), where identification of enriched repeat motifs is followed by ad-hoc realignment of relevant reads to the reference genome and manual evaluation of loci where these reads align.

EHdn has some limitations and areas for further improvement. It is limited to the detection of repetitive sequence longer than the read length and cannot, in general, detect shorter expansions. However, detection of these shorter expansions are feasible with the existing catalog-based methods, or SV detection methods. It may be possible to extend the detection limit to shorter repeat expansions, however increasing the search space will lead to increased runtime and reduced power to detect outlier expansions.

In many previous studies, identification of pathogenic REs required years of work and involved linkage studies to isolate the region of interest followed by targeted sequencing to identify the likely causative mutations. EHdn can be used as a front-line tool in such studies to rapidly identify candidate REs. The loci flagged by EHdn can be further studied by defining custom input files describing these novel REs for analysis with targeted methods, as well as other molecular assays. The benefits of this approach were demonstrated in a recent study, where EHdn successfully identified a novel complex pathogenic RE (Rafehi et al. 2019).

## Conclusions

We presented ExpansionHunter Denovo, a new genome-wide and catalog-free method to search for REs in WGS data. We demonstrated that EHdn consistently detects REs in real and simulated data. Given the widespread adoption of WGS for rare disease diagnosis, we expect that EHdn will enable further RE discoveries that will likely resolve the genetic cause of disease in many individuals.

## Methods

### Identification of IRRs

To determine if a read *r* is an in-repeat read, we first check the read for periodicity. We define *I*_*k*_ (*i*) = 1 if *r*_*i*_ = *r*_*i*+*k*_ and 0 otherwise, where *r*_*i*_ and *r*_*i*+*k*_ are the *i* th and (*i* + *k*) th bases of the read *r*. We then let 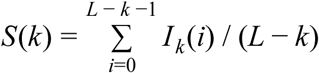 where *L* is the read length. Note that if a read consists of a perfect stretch of repeat units of length *k* then *S* (*k*) = 1. We search across of range of motif lengths (by default *k* ∈ {2, 3, …, 20}) for the smallest *k* such that *S* (*k*) ≥ *t* where *t* is a set threshold (we use *t* = 0.8 in all our analyses). If such a value of *k* is found, we extract the putative repeat unit using the most frequent bases at each offset 0 ≤ *i* ≤ *k* − 1. Since the orientation of the repeat where a given IRR originated is unknown in general, the unit of the repeat is ambiguous. To remove the ambiguity, we select the smallest repeat unit in lexicographical order under circular permutation and reverse complement operations. We then use this putative repeat unit to calculate a weighted-purity (WP) score of a read (Dolzhenko et al. 2017). We assume that a read is an IRR if it achieves WP score of at least 0.9. The WP score lowers the penalty for low-quality mismatches in order to account for the possibility of an increased base-call error rate that may occur in highly repetitive regions of the genome.

EHdn searches for IRRs among unaligned reads and reads whose mapping quality (MAPQ) is below a set threshold which in the analysis presented here was set to 40. For this study, we limited our analysis to motif lengths between two and 20 base pairs. Motif lengths equal to one were excluded to eliminate the large number of homopolymer repeats from the downstream analyses since we identified over 30 times as many homopolymer IRRs as IRRs with longer repeat motifs.

EHdn designates a read pair as a paired IRR if both mates are IRRs with the same repeat motif. A read is designated as an anchored IRR if it is an IRR and its mate is not an IRR and has MAPQ above a set threshold which was set to 50 for this study. Parameters such as the maximum allowed MAPQ for an IRR, the minimum allowed MAPQ for an anchor, and the range of repeat unit lengths for which to search are all tunable with EHdn. For example, setting the anchor read MAPQ threshold to 0 and the IRR MAPQ threshold to 60 ensures that every read pair in the alignment file is analyzed (assuming that the MAPQ values range from 0 to 60) at the cost of a corresponding increase in runtime.

### Merging IRRs

Because an anchored IRR is assigned to the location of the aligned anchor read and not the position of the actual repeat (whose exact location may be unknown), a single repeat may produce anchored IRRs at a variety of locations centered around the repeat. To account for this, anchored IRRs with the same repeat motif are merged if their anchors are aligned within 500 bp of one another. When multiple samples are analyzed, the anchor regions are also merged across all samples and the counts of anchored IRRs (normalized to 40x read depth) are tabulated for each merged region and sample. Additionally, the depth-normalized counts of paired IRRs are tabulated for each repeat motif and sample.

### Prioritization of expanded repeats

EHdn supports case-control and outlier analyses of the underlying dataset. The case-control analysis is based on a one-sided Wilcoxon rank-sum test. It is appropriate for situations where a significant subset of cases is expected to contain expansions of the same repeat.

The outlier analysis is appropriate for heterogeneous cohorts where enrichment for any specific expansion is not expected. The outlier analysis bootstraps the sampling distribution of the 95% quantile and then calculates the z-scores for cases that exceed the mean of this distribution. The z-scores are used for ranking the repeat regions. Similar outlier-detection frameworks were also developed for exSTRa (Tankard et al. 2018) and STRetch (Dashnow et al. 2018).

Both the case-control and the outlier analyses can be applied either to the counts of anchored IRRs or to the counts of paired IRRs. We refer to these as locus or motif methods, respectively. The high-ranking regions flagged by the analysis of anchored IRRs correspond to approximate locations of putative repeat expansions. The high-ranking motifs flagged by the analysis of paired IRRs correspond to the overall enrichment for repeats with that motif.

### Defining relevant repeat expansions

A catalogue of pathogenic or potentially pathogenic repeat expansions was collated from the literature. We supplemented this catalog with recently reported STRs linked with gene expression (Fotsing et al. 2019), and repeats with longer motifs overlapping with disease genes (Supplemental Information).

### Simulated repeat expansions

Expanded repeats were simulated using a strategy similar to that taken by BamSurgeon (Ewing et al. 2015). Briefly, we simulated reads in a 2Kb region around an expanded repeat and then aligned the reads to the reference genome. We then removed reads in the same region from a WGS control sample and merged alignments of real and simulated data together (Figure S2; Supplemental Information).

### Human WGS data

The control WGS samples are from Illumina Polaris dataset (Illumina n.d.). All samples were sequenced on an Illumina HiSeqX instrument using TruSeq DNA PCR-free sample prep. The 91 Coriell samples with experimentally-confirmed repeat expansions in *DMPK, FMR1, FXN* and *HTT* were introduced in our earlier publication (Dolzhenko et al. 2017).

## Funding

BT was funded by the Canadian Institutes for Health Research Banting Postdoctoral Fellowship and the Canadian Open Neuroscience Platform Research Scholar Award. MB was supported by an Australian National Health and Medical Research Council (NHMRC) Program Grant (GNT1054618) and an NHMRC Senior Research Fellowship (GNT1102971). This work was made possible through Victorian State Government Operational Infrastructure Support and Australian Government NHMRC IRIISS.

## Notes

https://github.com/Illumina/ExpansionHunterDenovo

