## Supplemental Information for "ExpansionHunter Denovo: A computational method for locating known and novel repeat expansions in short-read sequencing data"

### Supplemental Methods

#### Initial processing of WGS data

Human and simulated WGS data was aligned to the GRCh37-lite genome

- http://www.bcgsc.ca/downloads/genomes/9606/hg19/1000genomes/bwa_ind/genome/GRCh37-lite.fa

with BWA mem v0.7.17 (Li 2013) using samtools v1.9 (Li et al. 2009). WGS data that was originally obtained in BAM format was converted to FASTQ using Bazam v1.0.1 (Sadedin and Oshlack 2019).

#### Detection of repeat expansions

ExpansionHunter Denovo v0.8.6 was used to generate the genome-wide STR profile for each WGS sample like so:

ExpansionHunterDenovo profile --reads SAMPLE.bam --reference GRCh37-lite.fa --output-prefix SAMPLE

The manifest file was synthesized for each required comparison, then multisample STR profiles generated and subsequent locus- and motif-based analyses were performed like so:

ExpansionHunterDenovo merge --reference GRCh37-lite.fa --manifest MANIFEST.tsv --output-prefix OUTPUT python3 casecontrol.py locus --manifest MANIFEST.tsv --multisample-profile OUTPUT.multisample_profile.json --output-prefix casecontrol_locus.tsv

python3 casecontrol.py motif --manifest MANIFEST.tsv --multisample-profile OUTPUT.multisample_profile.json --output-prefix casecontrol_motif.tsv

python3 outlier.py locus --manifest MANIFEST.tsv --multisample-profile OUTPUT.multisample_profile.json --output-prefix outlier_locus.tsv

python3 outlier.py motif --manifest MANIFEST.tsv --multisample-profile OUTPUT.multisample_profile.json --output-prefix outlier_motif.tsv

The output of each command was sorted by p-value/z-score and the rank of each relevant locus or motif was extracted to evaluate performance.

STRetch (https://github.com/Oshlack/STRetch, commit 5405902) was run using the recommended pipeline for WGS analysis starting from BAM files for each sample independently (STRetch_wgs_bam_pipeline.groovy). The STRetch STR catalog was converted to GRCh37 (sed s/chr// hg19.simpleRepeat_period1-6.dedup.sorted.bed > grch37_input_regions.bed) and set as the input_regions parameter. Each sample was compared to the included controls (hg19.PCRfreeWGS_143_STRetch_controls.tsv). The rank of each relevant repeat was extracted from the sorted STRs.tsv output file.

#### Baseline simulations

The 2x150bp reads were simulated using wgsim (lh n.d.) read simulator. The base error rate was set to 0, indel fraction to 0, and fragment length to 450 bp with a standard deviation of 50. Simulated reads were mapped to the reference genome as described above, and IRR pairs were identified from the BAM file using EHdn profile.

#### Comparison between STRetch and GangSTR STR databases

GangSTR and STRetch catalogs were downloaded on February 20th, 2019 from

- https://s3.amazonaws.com/gangstr/hg19/genomewide/hg19_ver13_1.bed.gz and
- https://figshare.com/s/1a39be9282c90c4860cd

respectively. These catalogs were compared with the list of pathogenic loci using the Intervene tool (Khan and Mathelier 2017) requiring at least 1 bp overlap. The comparison script, RunComparison.sh, is located here:

- https://github.com/egor-dolzhenko/ehdn-paper-analysis/tree/master/CompareSTRDatabases/

#### Simulation of repeat expansions

The repeat expansions were simulated following a strategy similar to the one used by BamSurgeon(Ewing et al. 2015). Briefly, a region around a non-expanded target repeat in a control WGS sample was replaced with synthetic reads supporting an expansion in a heterozygous (one expanded allele and one reference-length allele) state (Figure S2). Specifically,

- The WGS sample HG03522 from the Polaris Kids cohort (Illumina n.d.) was used as the control.
- A FASTA file containing the expanded repeat along with 2Kb flanking sequence upstream and downstream of the repeat was generated for read simulation.
- Reads were simulated from the FASTA file using ART v2.5.8 (Huang et al. 2012) at 18x allele read depth. ART parameters for read length (150bp), mean insert size (460), and insert size standard deviation (115) were chosen to match the control WGS sample.

- Processed simulated reads, same as described above, were merged with sample HG03522 using samtools (v1.3.1).

Further details about the simulation process can be found here:

- https://github.com/egor-dolzhenko/ehdn-paper-analysis/tree/master/STR_Simulation

#### Manual cataloging of repeats

Pathogenic repeats were catalogued from a variety of sources including recent reviews and original discovery publications with source citations listed (Table S1). Genomic coordinates of pathogenic repeats were manually validated using IGV (Robinson et al. 2011).

Twelve STRs with varying motif lengths were selected from a recently-described set of expression-linked STRs (Fotsing et al. 2019) (Table S2).

A set of 27 STRs with motifs of size 7-10bp were selected from the GangSTR catalog. They were chosen to overlap intronic or UTR regions of autosomal recessive genes from the OMIM catalog (“OMIM - Online Mendelian Inheritance in Man” n.d.) (Table S2).

### Supplemental Results

#### Additional simulation results

Since we anticipate that EHdn will be used for discovery of novel pathogenic REs, we additionally simulated expansions of STRs with an associated effect on gene expression(Fotsing et al. 2019). From the catalog of fine-mapped eSTRs, we selected 12 loci with varying motifs linked with known disease genes. Expansions with 200 copies were simulated at these loci and EHdn prioritized all 12 REs in the top five. STRetch also performed well at these loci, although missed three because they were not present in its catalog.

To highlight that EHdn is not limited to short STR motifs, we tested its capacity to detect an expansion of a known pathogenic repeat with 12 bp motif in the promoter region of the *CSTB* gene. Using EHdn we detected this simulated expansion at the pathogenic lower bound (40 copies) (Table S6). We further demonstrate the ability of our method to detect REs with longer motifs at other similar loci. We simulated expansions of 27 repeats with 7-10 bp motifs within genes implicated in autosomal recessive genetic diseases (Supplemental Methods). All 27 loci were ranked in the top five in both the locus and motif analyses (Table S6).

### Supplemental Figures

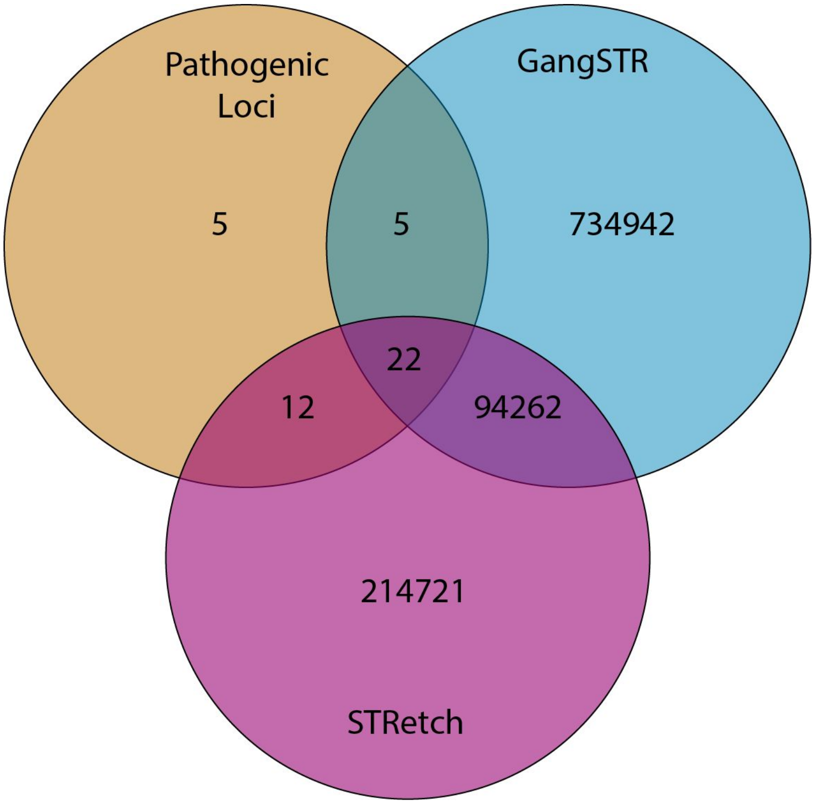


Figure S1 - Summary of concordance between STR catalogs: STRetch catalog (based on the UCSC simple repeats track), the GangSTR catalog, and a curated list of pathogenic or potentially pathogenic repeats reported in the literature.

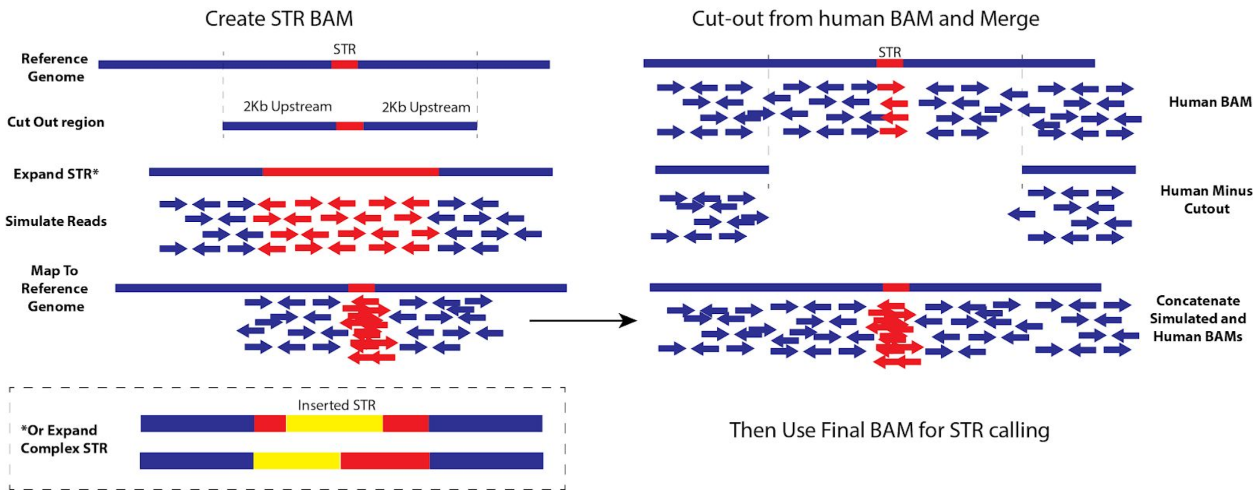


Figure S2: - Overview of simulating samples with repeat expansions. STR simulation by replacing reads around a non-expanded repeat in a control WGS sample with synthetic reads supporting an expansion.

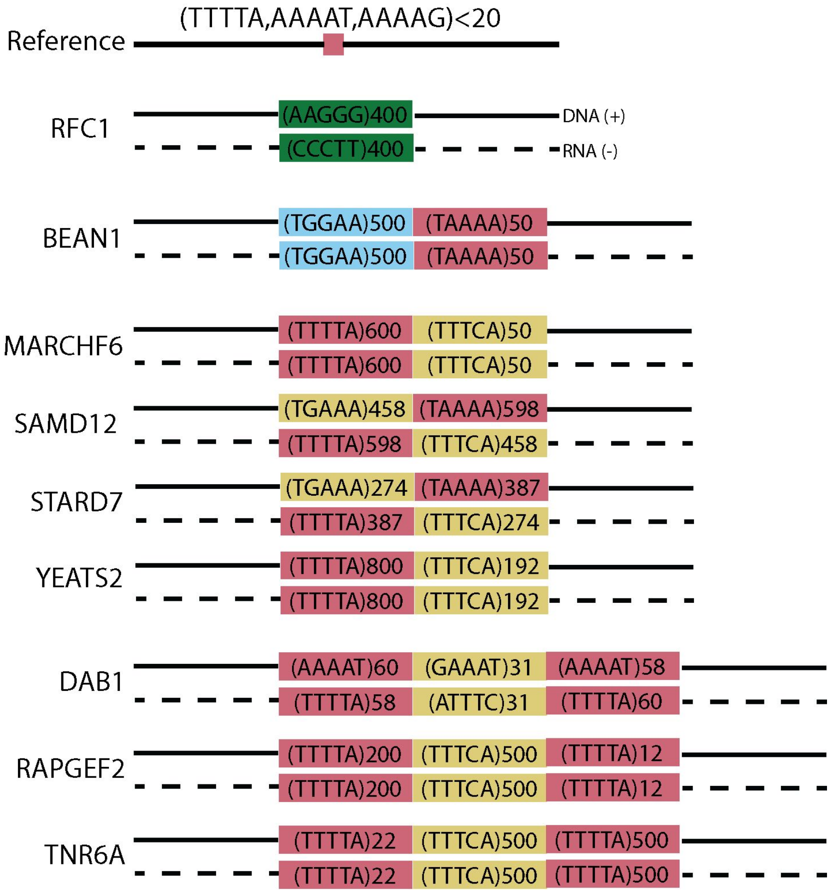


Figure S3 - Structure of nine complex pathogenic repeats.

### Supplemental Tables

Table S1
Definitions of small, large, degenerate, and complex repeat expansions. Gene, publication source, GRCh37 and GRCh38 reference coordinates in BED format (0-based half-open), repeat motif, pathogenic lower bound, motif in frame of gene, and presence in STRetch/GangSTR databases is listed for each repeat locus. For degenerate repeats, ambiguous nucleotides are represented with 'N' and the locus is defined with respect to the amino-acid expansion (e.g. poly-Alanine).

Table S2
Repeats with long motifs (motif length 7-10bp) and repeat loci linked to gene expression which were used for simulation. Associated gene, genic region (e.g. intronic, upstream, UTR5), GRCh37 coordinates, repeat motif, and simulated size in motif counts are provided for each locus.

Table S3
Results for simulations of 12 small pathogenic repeat expansions for STRetch, EHdn Locus, and EHdn Motif outlier prioritization analyses. STRs with variable nucleotides are omitted. Each row describes a single simulation of a repeat expansion, with the gene, pathogenic lower bound, motif, motif length, simulated size in copies, simulated size in base pairs, the STR tool which was used, the rank, the STR rank (only comparing motifs length 2-6bp), and the Z-score or p-value are listed. Missing values are represented by -1.

Table S4
Results for simulations of 22 large pathogenic repeat expansions for STRetch, EHdn Locus, and EHdn Motif outlier prioritization analyses. STRs with variable nucleotides are omitted. Columns shown are described in Table S3. Missing values are represented by -1.

Table S5
Results for simulations of 27 large-motif repeat expansions for EHdn Locus and EHdn Motif outlier prioritization analyses. Each repeat consists of 100 motif copies (motif

length 7-10bp). Both EHdn motif and EHdn locus prioritization results are shown. Columns shown are described in Table S3. Missing values are represented by -1.

Table S6
Results for simulations of small-motif expression-linked repeat expansions. Each repeat consists of 200 motif copies (motif length 2-6 bp). STRetch, EHdn motif, and EHdn locus prioritization results are shown. Columns shown are described in Table S3. Missing values are represented by -1.

Table S7
Results for simulations of nine complex pathogenic repeat expansions with EHdn Locus and EHdn Motif outlier prioritization analyses. Each locus is delineated by gene name, such that a single simulation is performed for each gene. Ranks for each motif present at a given locus are shown as separate rows annotated by sub-motif. Ranks are additionally listed for both outlier analysis performed with the parameter defining the maximum mapping quality threshold for IRRs (--max-irr-mapq) set to 40 (default) and 60. Missing values are represented by ‘N/A’.

Ewing, Adam D., Kathleen E. Houlahan, Yin Hu, Kyle Ellrott, Cristian Caloian, Takafumi N. Yamaguchi, J. Christopher Bare, et al. 2015. “Combining Tumor Genome Simulation with Crowdsourcing to Benchmark Somatic Single-Nucleotide-Variant Detection.” *Nature Methods* 12 (7): 623–30.

Fotsing, Stephanie Feupe, Jonathan Margoliash, Catherine Wang, Shubham Saini, Richard Yanicky, Sharona Shleizer-Burko, Alon Goren, and Melissa Gymrek. 2019. “The Impact of Short Tandem Repeat Variation on Gene Expression.” *Nature Genetics* 51 (11): 1652–59.

Huang, Weichun, Leping Li, Jason R. Myers, and Gabor T. Marth. 2012. “ART: A next-Generation Sequencing Read Simulator.” *Bioinformatics* 28 (4): 593–94.

Illumina. n.d. “Illumina/Polaris.” GitHub. Accessed December 1, 2019. https://github.com/Illumina/Polaris.

Khan, Aziz, and Anthony Mathelier. 2017. “Intervene: A Tool for Intersection and Visualization of Multiple Gene or Genomic Region Sets.” *BMC Bioinformatics* 18 (1): 287.

lh. n.d. “lh3/wgsim.” GitHub. Accessed November 29, 2019. https://github.com/lh3/wgsim.

Li, Heng. 2013. “Aligning Sequence Reads, Clone Sequences and Assembly Contigs with BWA-MEM.” *arXiv [q-bio.GN]*. arXiv. http://arxiv.org/abs/1303.3997.

Li, Heng, Bob Handsaker, Alec Wysoker, Tim Fennell, Jue Ruan, Nils Homer, Gabor Marth, Goncalo Abecasis, Richard Durbin, and 1000 Genome Project Data Processing Subgroup. 2009. “The Sequence Alignment/Map Format and SAMtools.” *Bioinformatics* 25 (16): 2078–79.

"OMIM - Online Mendelian Inheritance in Man." n.d. Accessed December 1, 2019. https://omim.org/.

Robinson, James T., Helga Thorvaldsdóttir, Wendy Winckler, Mitchell Guttman, Eric S. Lander, Gad Getz, and Jill P. Mesirov. 2011. "Integrative Genomics Viewer." *Nature Biotechnology* 29 (1): 24–26.

Sadedin, Simon P., and Alicia Oshlack. 2019. "Bazam: A Rapid Method for Read Extraction and Realignment of High-Throughput Sequencing Data." *Genome Biology* 20 (1): 78.
